# Wild-type MECP2 expression coincides with age-dependent sensory phenotypes in a female mouse model for Rett syndrome

**DOI:** 10.1101/2022.04.25.482695

**Authors:** Michael Mykins, Dana Layo-Carris, Logan Reid Dunn, David Wilson Skinner, Alexandra Hart McBryar, Sarah Perez, Trinity Rose Shultz, Andrew Willems, Billy You Bun Lau, Tian Hong, Keerthi Krishnan

**Affiliations:** Department of Biochemistry & Cellular and Molecular Biology, University of Tennessee, Knoxville, TN

## Abstract

Rett syndrome is characterized by an early period of typical development and then, regression of learned motor and speech skills in girls. Loss of MECP2 protein is thought to cause Rett syndrome phenotypes. The specific underlying mechanisms from typical developmental trajectory to regression features throughout life are unclear. Lack of established timelines to study the molecular, cellular, and behavioral features of regression in female mouse models is a major contributing factor. Due to random X-chromosome inactivation, female patients with Rett syndrome and female mouse models for Rett syndrome (*Mecp2^Heterozygous^*, Het) express a functional copy of wild-type MECP2 protein in approximately half of all cells. As MECP2 expression is regulated during early postnatal development and experience, we characterized the expression of wild-type MECP2 in the primary somatosensory cortex of female Het mice. Here, we report increased MECP2 levels in non-parvalbumin-positive neurons of 6-week-old adolescent Het relative to age-matched wild-type controls, while also displaying similar levels of perineuronal net expression, mild tactile sensory perception deficits, and efficient pup retrieval behavior. In contrast, 12-week-old adult Het express MECP2 at levels similar to age-matched wild-type mice, show increased perineuronal net expression in the cortex, and display significant tactile sensory perception deficits. Thus, we have identified a set of behavioral metrics and the cellular substrates to study regression during a specific time in the female Het mouse model, which coincide with changes in wild-type MECP2 expression. We speculate that the precocious increase in MECP2 expression within specific cell types of adolescent Het may provide compensatory benefits at the behavioral level, while the inability to further increase MECP2 levels leads to regressive behavioral phenotypes over time.

## Introduction

Rett syndrome (RTT) is a neuropsychiatric disorder, predominantly caused by mutations in the X-linked gene Methyl CpG-Binding Protein 2 *(MECP2)* (Amir et al., 1999; Hagberg et al., 1983; Rett, 1966). Most surviving patients with RTT are females who are heterozygous for *MECP2* mutations. Diagnostic criteria for RTT include partial or complete loss of acquired purposeful motor skills or spoken language during a regression period in the first few years of life. Additionally, primary sensory processing is disrupted in RTT patients (Buchanan et al., 2019; Djukic et al., 2012; Djukic and Valicenti McDermott, 2012; Key et al., 2019; LeBlanc et al., 2015; Merbler et al., 2020; Nomura, 2005; Peters et al., 2015; Stallworth et al., 2019; Symons et al., 2019). Such disruptions during development and adulthood contribute to motor dysfunction and social communication deficits in girls and women with RTT. Patients typically survive into middle age with periods of plateau and regression throughout life (Bebbington et al., 2008; Buchanan et al., 2019; Charman et al., 2002; Djukic et al., 2012; Djukic and Valicenti McDermott, 2012; Einspieler and Marschik, 2019; Kerr, 1995; Kirby et al., 2010; Nomura, 2005). Rodent models recapitulate many of the features of RTT, including sensory processing, social communication deficits, and motor difficulties (Durand et al., 2012; Ito-Ishida et al., 2015; Krishnan et al., 2015; Krishnan et al., 2017; Lau et al., 2020a; Lee et al., 2017; Lo et al., 2016; Orefice et al., 2016). Though much is known about the characteristics of RTT phenotypes, the underlying cause for regression during various stages of life and the changes in syndromic phenotypes over time is unclear. The prevailing idea is that loss of MECP2 expression and function is the major driver for RTT phenotypes; however, the role of the wild-type MECP2 protein in half of the cells in RTT is unknown.

In RTT, random X-chromosome inactivation leads to mosaic wild-type MECP2 expression in approximately 50% of most major cell types in the body, including neurons (Samaco et al., 2013; Young and Zoghbi, 2004). Due to this inherent mosaicism, individual girls and women with RTT, as well as individual female heterozygous mice *(Mecp2^heterozygous^*, Het), are unique. However, the impact of mosaic MECP2 expression on cellular function in Het is not fully understood. We hypothesize that the mosaicism in Het contributes to the observed syndromic phenotypes, due to likely compensation and adaptations in individual cells throughout life. In rodents, MECP2 expression increases postnatally, concomitant with the emergence of appropriate critical periods, and is generally thought to be stable in adolescence and adulthood (Braunschweig et al., 2004; Skene et al., 2010; Yagasaki et al., 2018). MECP2 is variably expressed across cell types, with higher expression in GABAergic interneurons in comparison to pyramidal neurons within the same region (Braunschweig et al., 2004; Ito-Ishida et al., 2015; Picard and Fagiolini, 2019; Skene et al., 2010; Takei et al., 2021; Yagasaki et al., 2018). Intriguingly, the status of wild-type MECP2 expression over time, in Het, is unknown. Due to its significant roles in gene regulation (Chahrour et al., 2008; Chen et al., 2015; Fuks et al., 2003; Gabel et al., 2015; Jones et al., 1998; Klose et al., 2005; Lyst and Bird, 2015; Tao et al., 2009; Zhou et al., 2006) and the timing of experience-dependent plasticity (Krishnan et al., 2015, 2017; Lau et al., 2020a; Morello et al., 2018; Noutel et al., 2011; Patrizi et al., 2020; Picard and Fagiolini, 2019), investigation of wild-type MECP2 expression in Het across ages is essential to unravel the mechanisms connecting genotype to phenotype throughout the lifetime of the individual.

Previous work found 6-week-old adolescent Het are thought to not exhibit overt symptoms and 12-week-old adult females are considered early post-symptomatic (Samaco et al., 2013). To specifically address social communication and complex sensorimotor integration, we have used a pup retrieval task, an ethologically-relevant behavior involving primary sensory processing and sensorimotor integration in an untrained task of retrieving scattered pups back to the nest (Beach and Jaynes, 1956; Lonstein et al., 2015; Stern, 1996). Adult Het are inefficient at the pup retrieval compared to age matched wild-type (WT) females (Krishnan et al., 2017; Stevenson et al., 2021). Furthermore, adult Het exhibit atypical tactile interactions with pups involving the whiskers and face, with decreased grooming behavior and increased proximal pup approaches (Stevenson et al., 2021). Together, these results suggest abnormal tactile processing in Het, which could contribute to their inefficient pup retrieval. Tactile processing is particularly relevant to alloparental care, as sensory cues from offspring are singularly essential for the early retention and maintenance of maternal behavior (Hinde and Davies, 1972; Lonstein and Stern, 1999; Maier, 1963; Morgan et al., 1992; Rosenblatt, 1967; Stern, 1996; Stern and Kolunie, 1989; Stern and Mackinnon, 1978). Thus, we focused on the tactile sensory perception modality in adolescent and adult Het using texture discrimination and object recognition tasks (Antunes and Biala, 2012; Diamond et al., 2008; Orefice et al., 2016, Orefice et al., 2019; Wu et al., 2013).

Here, we report the first evidence of a precocious increase of wild-type MECP2 expression within MECP2-expressing cells 6-week-old adolescent Het relative to age-matched WT; these MECP2 expression changes are observed in non-parvalbumin positive (PV+) GABAergic neurons, which exhibit lower levels of MECP2, but not in PV+ GABAergic neurons exhibiting higher MECP2 levels. This cell-type specific expression difference is not maintained in 12-week-old adult Het, suggesting that cells expressing wild-type MECP2 protein in Het are regulated by age-specific mechanisms. This MECP2 expression phenotype in adolescent Het is concomitant with typical structural plasticity, as measured by the expression of specialized extracellular matrix structures ensheathing PV neurons called perineuronal nets (PNNs). Furthermore, a systematic, frame-by-frame analyses of texture and object investigation tasks in conjunction with pup retrieval behavior reveal that adolescent Het behave similarly to adolescent WT in both simple tactile and complex social behaviors. Adult Het mice, however, are known to be inefficient at pup retrieval (Krishnan et al., 2017; Stevenson et al., 2021) and display overt tactile sensory perception issues, suggesting that observed tactile hyposensitivity in Het is age-related. Based on these findings, we hypothesize that precocious increase of wild-type MECP2 expression in specific cell types of adolescent Het provides compensatory benefits at the behavioral level, while the inability to further increase MECP2 levels with age leads to observed RTT stereotypies over time. Together, these novel results open new directions of exploration on the long-term, cell-type specific roles of MECP2 expression, regulation, and function in Het while emphasizing the need to analyze dynamics of cellular and behavioral phenotypes.

## Materials and Methods

### Animals

We used the following mouse strains: CBA/CaJ (JAX:000654) and *Mecp2^heterozygaus^* (Het) (C57BL/6J, *B6.129P2(C)-Mecp2^tm1·1Bird^*/J, JAX:003890) (Guy et al., 2001). All animals were group-housed by sex after weaning, raised on a 12/12-hour light/dark cycle (lights on at 7 am) and received food and water ad libitum. Behavioral experiments were performed using Het and female *Mecp2^WT^* siblings (WT) at either 6 (adolescent) or 10-to-12 (adult) weeks of age between the hours of 9 am and 6 pm (during the light cycle). All procedures were conducted in accordance with the National Institutes of Health’s Guide for the Care and Use of Laboratory Animals and approved by the Institutional Animal Care and Use Committee at the University of Tennessee-Knoxville.

### Immunohistochemistry

All animals were perfused with 4% paraformaldehyde (PFA) in 1X PBS immediately after behavior. Extracted brains were stored in 4% PFA overnight at 4°C before storing long-term in PBS. Tissues were processed in cohorts (one WT and one Het) as previously described (Lau et al., 2020b). Briefly, coronal brain sections (100 μm thickness) encompassing the entire primary somatosensory cortex (SS1) were collected using a freezing microtome. We denoted the left hemisphere by poking a small hole in the ventral part of the brain. Free-floating sections were blocked with a 10% normal goat serum and 0.5% Triton X-100 in 1X PBS. Blocking solution at half the original concentration was used for antibody incubations. Tissues were incubated with biotin-conjugated WFA lectin (1:500; Sigma-Aldrich), MECP2 primary antibody (1:1,000; Cell Signaling), and PV antibody (1:1,000; Sigma-Aldrich) followed by secondary antibodies conjugated with AlexaFluor-488, Texas-Red, and AlexaFluor-405 (1:1,000; Invitrogen) to label PNNs, MECP2, and PV (if applicable), respectively. Tissues were counterstained with DAPI before mounting section onto slides.

### Microscopy and Image Analysis

#### Image Acquisition

*MECP2*: Image acquisition of both MECP2 and DAPI signals were performed using a VT-Hawk 2D-array scanning confocal microscope (Visitech intl.) mounted to an IX-83 inverted optical microscope (Olympus). Z-series images with a step interval of 0.5 μm were collected using a 60X objective lens (Nikon) in MetaMorph (Molecular Devices). Settings for each laser channel were determined in a similar manner as previously described for PNNs while utilizing the histogram function in MetaMorph. Acquisition settings were determined using the top 25.0 μm of WT tissues, and these settings were used to image all conditions in the 16-bit range. PV and PNN images (Figure 1a) were acquired in the same manner using a 40X objective lens (Nikon). Images were collected from the Layer IV barrel field subregion (S1BF) in both hemispheres of two sections per brain.

**Figure 1.**
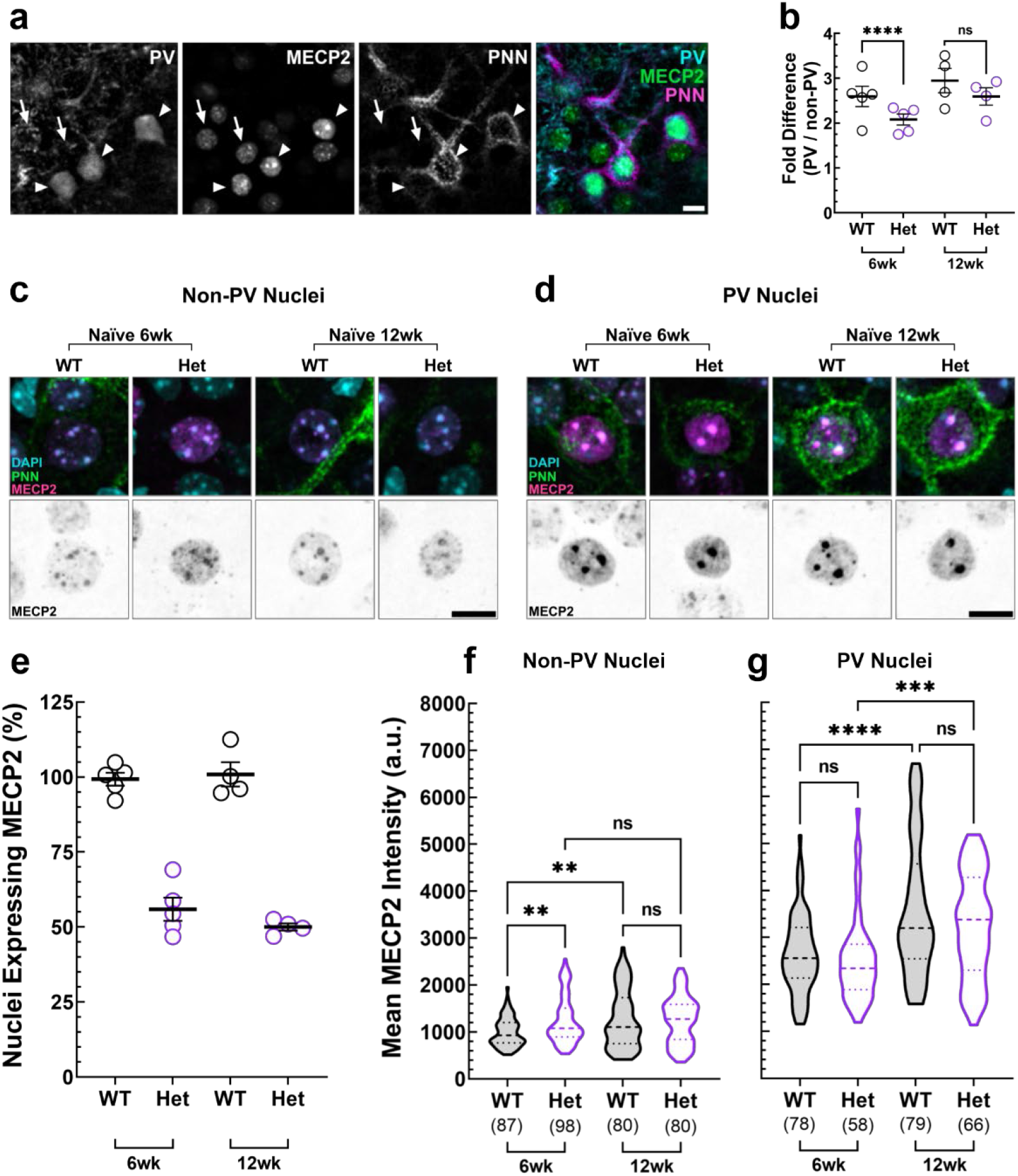
MECP2 is expressed differentially between cell types in S1BF of adolescent and adult WT and Het. **(a)** Representative 40X confocal images from Layer IV S1BF showing PNN colocalization with PV in MECP2+ nuclei. Arrows (non-PV) and arrowheads (PV) point to representative nuclei for quantification. Scale bar = 10 μm. **(b)** Mean intensity of MECP2 protein from PV and non-PV nuclei is represented as a fold difference (PV:non-PV ratio; number of replicates same as reported for figure 1f). **(c, d)** Representative nuclei stained with DAPI and MECP2 from non-PV (PNN-negative cells) (c) and PNN-positive parvalbumin (PV) interneurons (d) in adolescent and adult naïve WT and Het show that MECP2 intensity is higher within PV nuclei across ages, and non-PV nuclei of adolescent (6wk) Het appear more intense compared to age-matched WT or adult (12wk) Het. Scale bars = 10 μm. Het express MECP2 in approximately 50% of nuclei, compared to WT across ages, as measured by percentage of MECP2-positive nuclei in adolescent and adult WT and Het (n = 2 images per hemisphere per brain). **(f-g)** Violin plots of the mean intensity of nuclear MECP2 within (f) non-PV and (g) PV nuclei. MECP2 expression increased significantly with age in non-PV nuclei of WT, while MECP2 expression in non-PV cells remained consistent across ages. Compared to age-matched WT control, 6-week-old Het showed a significant increase in MECP2 expression in non-PV cells, while 12-week-old Het did not. In PV nuclei, MECP2 expression increased significantly with age within both genotypes; there were no significant differences between WT and Het MECP2 expression at either age. Respective ages are indicated on the x-axes, and the number of nuclei from which data was collected is shown in parentheses for each condition (n = 5 animals per genotype at adolescent age; n = 4 animals per genotype at adulthood). *Kruskal-Wallis followed by Uncorrected Dunn’s test: ns = not significant; **p < 0.01, ***p < 0.001.* **(b, e)** Mean ± SEM are shown.

*PNN:* Single-plane images of coronal tissue sections were collected using a Keyence epifluorescence microscope (Keyence BZ-X710) with a 10X objective and stitched using the BZ-X Analyzer software. We determined exposure settings for each cohort based on Het PNN intensity, as we previously showed atypical increased expression of PNNs in adult Het (Lau et al, 2020b). To reduce overexposure, we set the exposure time to the point when the first saturated pixel appeared in the sample, then reduced the exposure time by one unit. This procedure was performed in both hemispheres for all Het sections, and the lowest exposure time occurring most frequently was used to image all sections within that cohort.

#### Image Processing & Analysis

*MECP2:* Nuclear expression of MECP2 was quantified using ImageJ (Schneider et al., 2012). For initial background subtraction, a duplicate of the original image was created and processed by applying a median filter with a 32-pixel radius. The filtered image was then subtracted from the original image using the Image Calculator tool to produce 16-bit images for quantification. Mean MECP2 intensity was quantified within individual nuclei by creating 3.0-5.0 μm maximum intensity projections (16-bit greyscale) within the top 25.0 μm of each tissue. Projections were created using the same image plane for both the DAPI and MECP2 channels and utilized for ROI creation via the Polygon drawing tool and quantification, respectively. DAPI signal, PNN presence or absence, and MECP2-expression status were identified for individual cells using merged tricolor images. Statistical analyses and figures were generated using GraphPad Prism. The percentage of nuclei expressing MECP2 was calculated using the EzColocalization ImageJ plugin (Stauffer et al., 2018). Briefly, the DAPI and MECP2 z-series image stacks were used for reporter channels 1 and 2, respectively, with default threshold algorithms applied. The mean Mander’s Colocalization Coefficient (MCC) for MECP2 colocalization with DAPI signal was collected across all imaging planes. This process was repeated for the four images collected for each animal (2 per hemisphere) and averaged to yield an animal-specific mean MCC value. To compare MCC values in a standardized manner, the average MCC value for all WT images across both ages was calculated. Then, the mean MCC value for each WT and Het replicate was divided by the WT average MCC and multiplied by 100 to approximate the percentage of nuclei expressing MECP2.

*PNN*: Images were mapped and regions-of-interests (ROIs) for S1BF were outlined by overlaying the images with coronal maps from Paxinos and Franklin’s mouse atlas (4^th^ edition, 2013). The density of high-intensity PNNs were quantified as previously described (Lau et al., 2020b) using ImageJ (Schneider et al., 2012). Briefly, high-intensity PNNs were quantified by maximizing the contrast of each image and manually marking the signals that were 80% or more of the original PNN. Density was calculated as total number of high-intensity PNNs divided by ROI area. We also analyzed the full spectrum of PNN expression in the mapped images. 8-bit histograms specific to the green channel (PNNs) were generated using the Histogram tool in ImageJ. We extracted the distributions of strong PNN expression from all images using two-component Gaussian mixture models (GMM; see Figure 2) (Bibbings et al., 2019). The means of these distributions (red traces in Figure 2d) were used as representative summary statistics in subsequent analyses. We calculated intraclass correlation coefficients (ICCs) to show that the ratio of inter-individual variance to the total variance was moderate (0.218), such that comparisons on genotypes can be made. We built a linear mixed-effects model to confirm that the inter-individual variability did alter the statistical significance (Yu et al., 2022). The increased variability in Het compared to WT is likely due to natural variation due to random X-chromosome inactivation patterns in Het.

**Figure 2.**
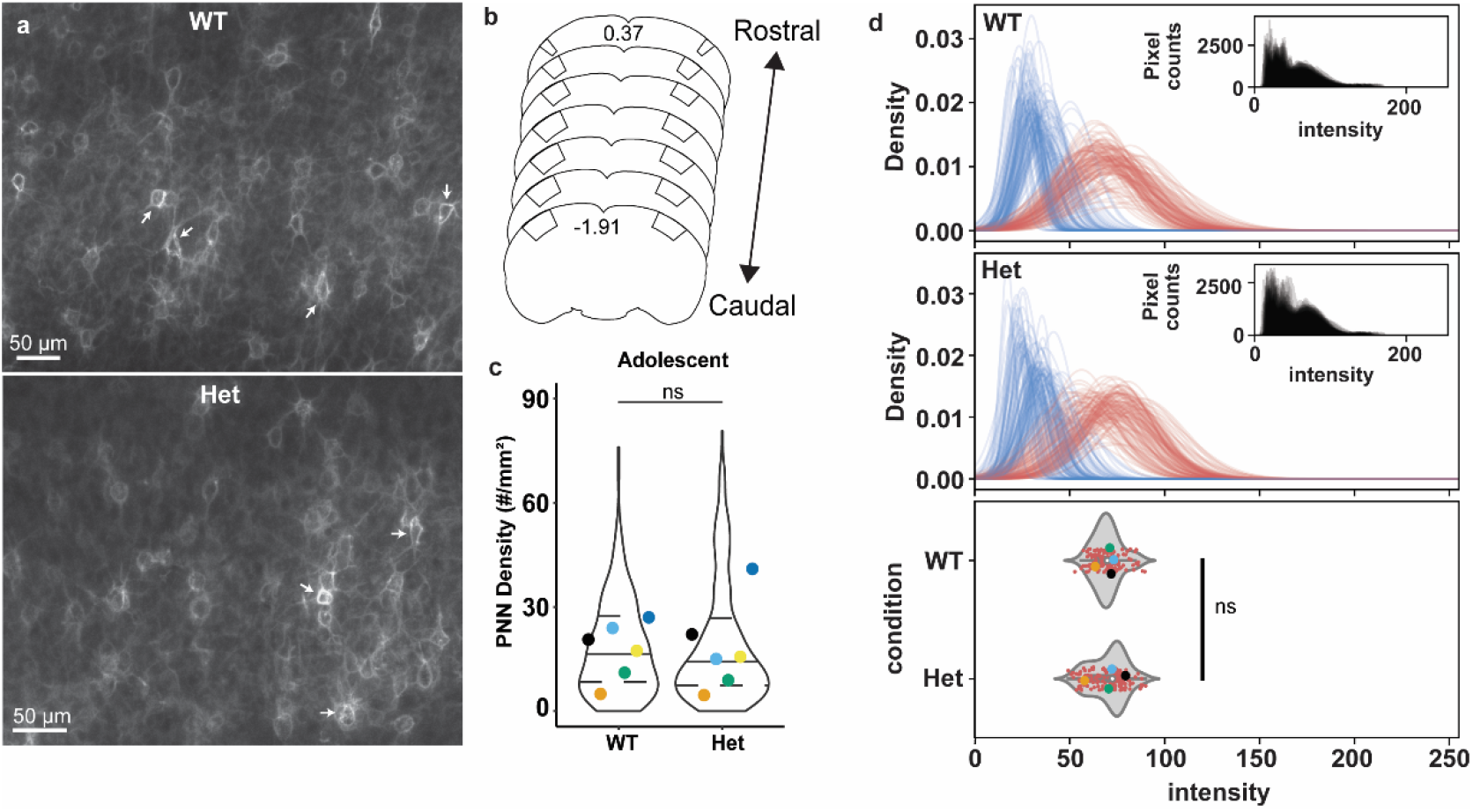
Adolescent Het show similar PNN density as WT within S1BF. **(a)** Representative epifluorescent 10x micrographs of PNNs in S1BF of adolescent WT and Het. Arrows indicate high-intensity PNNs quantified for analysis. **(b)** Schema of coronal mouse brain sections, depicting rostral (Bregma: 0.37 mm) to caudal (Bregma: −1.91 mm) regions of S1BF (boxes) analyzed for this study. Coordinate maps are based on Paxinos and Franklin’s mouse brain atlas, 4th edition. **(c)** Statistical analysis of manually counted high-intensity PNNs revealed that the density is similar between adolescent WT and Het. Each circle represents the average high-intensity PNN density of each brain, superimposed on violin plots showing median with distirbution of all sections. Individual colors represent cohorts processed and analyzed together. WT: n=307 images, 6 animals. Het: n=313 images, 6 animals. *Mann-Whitney test: ns = not significant.* **(d)** All intensity analysis of PNN expression. The top two panels show two distributions of PNN intensity obtained with a two-component Gaussian mixture model for each image (see **Methods**). The distributions with higher means (red traces) were used for subsequent analyses, while background signal (blue traces) was excluded. Each trace represents PNN intensity distribution of an image. Insets show histograms of raw intensities for pixels of all images (135 WT and 140 Het). Bottom panel: statistical analysis revealed no significant difference in all PNN intensity between adolescent WT and Het. The graph shows the distribution of the means (small red dots and violin plots) of the red traces in top panels. Large dots show aggregated mean of the red distributions for each color-coded cohort. Box plots within violin plots show 25^th^, 50^th^ and 75^th^ quartiles. *Linear mixed effect model and t-test: p > 0.05.*

### Pup Retrieval Task

#### Behavioral Paradigm

The pup retrieval task was performed as previously described (Stevenson et al., 2021). Two 4.5-week-old naïve females (one WT and one Het) with no prior experience of pup retrieval were co-housed with a pregnant CBA/CaJ female (10-12 weeks of age) 3-5 days before birth; naïve females were termed ‘surrogates’ upon cohousing. Once the pups were born (postnatal day 0; P0), we performed the pup retrieval task with the surrogates for six consecutive days. The behavior was performed in the home cage placed inside of a sound- and light-proof box as follows: one surrogate was placed in the home cage with 3-5 pups *(habituation;* 5 minutes), the pups were removed from the nest *(isolation;* 2 minutes), and the pups were scattered throughout the home cage by placing them in the corners and center, allowing the surrogate to retrieve pups back to the nest *(retrieval;* 5 minutes). The nest was left empty if there were fewer than five pups. The assay was performed in the dark and recorded using infrared cameras during all phases of each trial. If all pups were not retrieved to the nest by the 5-minute mark of the retrieval phase, we placed pups back in the nest in preparation for the next surrogate. After the behavior was completed, all mice were returned to the home cage.

#### Behavior Analysis

All videos were coded so the analyzers were blind to the identity of the mice. Each video was manually scored using a latency index (the amount of time to retrieve all pups back to the nest out of 5 minutes) and the number of errors (surrogate-pup interactions that did not result in successful retrieval). Latency index was calculated as:

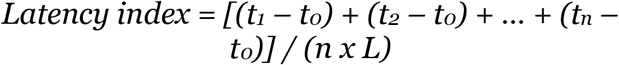

Where *n* = number of pups outside the nest, *t*_o_ = start time of trial (s), *t*_n_ = time of the nth pup’s successful retrieval to the nest (s), *L* = total trial length (300 s). Statistical analyses and figures were generated using GraphPad Prism.

#### Tactile Perception Tasks

WT and Het underwent a five-day battery of behavioral tests to assay tactile processing through the whiskers (Figure 4). The behavior chamber used for tactile assays was a 40 cm x 40 cm x 40 cm plexiglass box with two sets of holes in the center of the plexiglass floor for the texture/object inserts. Holes were drilled to be equidistant from the center, with a 4.0 cm spacing between hole sets. Texture discrimination plexiglass inserts were 0.5 cm (thickness) x 4.0 cm (width) x 15.0 cm (height). Rodents use their whiskers to both detect and discriminate between textures and objects (Rodgers et al., 2021). We chose to test the ability of experimental mice to discriminate between textures using both large and small textural differences. Sandpaper textures were designed to slide on and off the inserts to be cleaned between animals. The textures were made of 3M sandpaper of varying grit: a course 60-grit, a medium 120-grit, and a fine 220-grit. Inserts used for novel object recognition were plexiglass spheres (4.0 cm diameter) and plexiglass cubes (64 cm^3^) (Orefice et al., 2016; Wu et al., 2013). The chamber and inserts were cleaned with MB-10 chlorine dioxide disinfectant and allowed to fully dry before the start of each behavior, and an additional 5-minute period was allotted between tests to eliminate scent biases. All behaviors were performed in the dark and recorded with an infrared camera.

**Figure 3.**
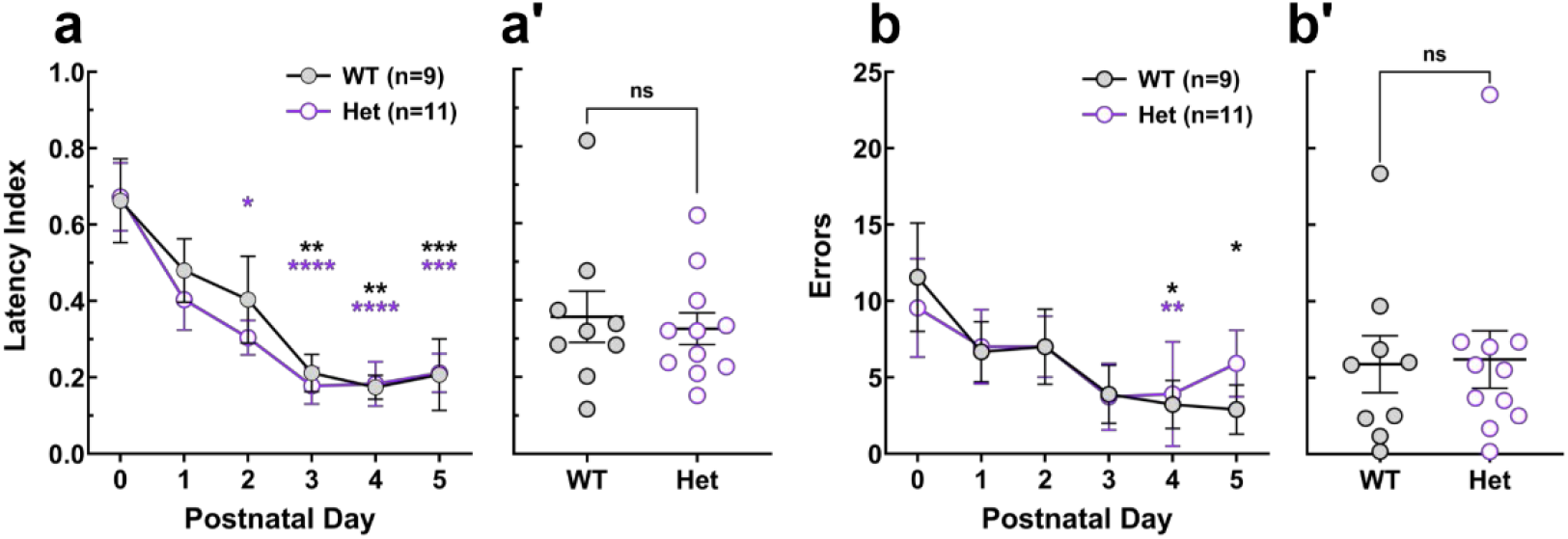
Adolescent Het perform pup retrieval comparable to WT. **(a-a’)** Mean latency index of pup retrieval behavior for each testing day (a) and averaged per animal across all testing days (a’) showed significant improved retrieval across testing days for both WT and Het, and no significant difference in average latency between genotypes. **(b-b’)** Mean number of errors during pup retrieval for each testing day (b) and averaged per animal across all testing days (b’) showed same result as latency index. **(all panels)** Mean ± SEM are shown. **(a, b)** *Kruskal-Wallis followed by Uncorrected Dunn’s test; *p < 0.05, **p < 0.01, ***p < 0.001, ****p < 0.0001.* Color * = within genotype, significance between P0 and P-day. **(a’, b’)** *Mann-Whitney test: ns = not significant.*

**Figure 4.**
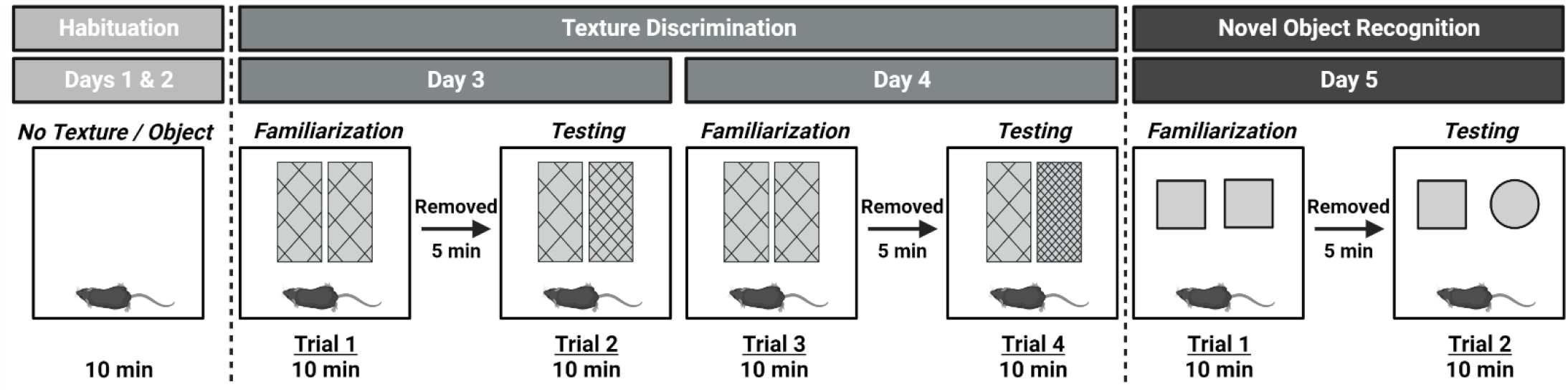
Tactile behavior schema. Behavior tests begin with two days of habituation to introduce the mice to the behavioral box set up. On each days of texture discrimination (days 3 and 4), individual mouse undergoes a 10-minute familiarization trial followed by a 10-minute testing trial, with varied sandpaper grits. On day 5, objects are used instead of textures for novel object recognition task. See Methods for detail.

#### Behavioral Paradigm

##### Habituation

On days 1 and 2, mice were placed individually into the behavior chamber for 10 minutes to habituate themselves with the new environment.

##### Texture Discrimination

On days 3 and 4, texture discrimination assays began with a familiarization trial. Experimental mice were individually placed into the chamber with two identical texture inserts and allowed to investigate for 1o minutes. Afterwards, the mouse was returned to the home cage for 5 minutes. During this time, textures and inserts for familiarization were replaced with new sets of inserts and textures for use in the testing trial. After 5 minutes, the mouse was returned to the behavior chamber for a second 10-minute trial (testing phase). Familiarization phases of each behavior were performed using two course texture inserts. During testing phase on days 3 and 4, we replaced one insert with a medium and fine texture insert, respectively.

##### Novel Object Recognition

On day 5, a novel object recognition assay was performed similar to the texture discrimination assay with minor changes. The familiarization trial was performed using two plexiglass cubes. For the testing trial, one plexiglass cube was replaced by a plexiglass sphere.

#### Behavior Analysis

Behaviors for both texture and object investigation were manually scored using Datavyu software (Datavyu Team, version 1.3.7, 2014). All analyzers were blind to the age and genotype of the mice. The videos were scored for the time intervals in which the mouse interacted with the novel and familiar textures/objects throughout the 10-minute testing trial. We classified interactions into three categories, as shown in Table 1, based on pilot observations and previous studies (Orefice et al., 2016; Pacchiarini et al., 2017; Wu et al., 2013). We excluded mice from analysis if the investigation time of textures/objects during a given trial did not exceed 2 seconds (Orefice et al., 2016; Patrick Wu and Dyck, 2018) (Table 2). After manual scoring, interaction sequences for each animal were extracted from Datavyu for quantitative analyses. We classified number of contacts, total investigation time, and duration per interactions with textures/objects into whisker, forepaw, and whole-body climbing. Our results are reported as raw number of contacts, raw total investigation times, and raw duration per interaction split by interaction type, age, and genotype. We conducted unpaired Mann Whitney-U tests to determine statistical significance between genotypes and across ages within the same genotype. Statistical analysis and data visualization were performed in R studio.

**Table 1.** Definitions of parameters used to score behaviors.

|  |  |
| --- | --- |
| Whisker | - Mouse within 3 cm of the object (due to typical whisker length) and actively exploring around texture/object |
| Whole-body climbing | - All four paws are off the ground and animal is climbing the texture/object |
| Forepaw | - Touching the texture/object with forepaws while hindlimbs are contacting the ground |

**Table 2.** Number of animals included in and excluded from analysis. Animals were excluded from analysis due to technical difficulties with video lagging/skipping or no interactions with textures.

| Age | Behavioral Test | Included |  | Excluded |  |
| --- | --- | --- | --- | --- | --- |
|  |  | WT | Het | WT | Het |
| 6WO | Texture Discrimination Day 1 (Trial 1 & 2) | 8 | 8 | 1 | 2 |
|  | Texture Discrimination Day 2 (Trial 3 & 4) | 9 | 10 | 1 | 0 |
|  | Novel Object Recognition (Trial 1 & 2) | 9 | 10 | 0 | 0 |
| 12W O | Texture Discrimination Day 1 (Trial 1 & 2) | 8 | 7 | 1 | 1 |
|  | Texture Discrimination Day 2 (Trial 3 & 4) | 9 | 8 | 1 | 0 |
|  | Novel Object Recognition (Trial 1 & 2) | 9 | 8 | 0 | 0 |

## Results

### Wild-type MECP2 expression is dynamically regulated in Het over age and cell types

MECP2 protein expression varies in different cortical cell types (Ito-Ishida et al., 2015; Takei et al., 2021). However, it is currently unknown if cell-type specific MECP2 expression is stable over age or if wild-type MECP2 expression in Het is similar to age-matched WT controls. Het expression of wild-type MECP2 was reported to be decreased relative to age-matched WT females; however, the brain region(s) and the specific ages of mice used in this investigation were not reported (Braunschweig et al., 2004). Thus, we performed immunofluorescent staining and confocal microscopy to first characterize MECP2 expression within individual nuclei of naïve WT and Het female mice during adolescence and adulthood. We chose to investigate expression within Layer IV of the barrel field subregion of the primary somatosensory cortex (S1BF), the cortical input layer from the whiskers, due to its importance in tactile sensory perception.

Previously, we reported that adult Het exhibit dysregulation of PV+ GABAergic neurons in the auditory and primary somatosensory cortices (Krishnan et al., 2017; Lau et al., 2020a; Lau et al., 2020b). Particularly, adult Het mice exhibit increased and atypical expression of perineuronal nets (PNNs) in the S1BF (Lau et al., 2020b), extracellular matrix structures mainly surrounding cortical PV+ GABAergic interneurons to structurally restrict plasticity (de Vivo et al., 2013; Hartig et al., 1992; Krishnan et al., 2017; Pizzorusso et al., 2002; Ueno et al., 2018). Thus, we focused MECP2 expression analysis in PNN+ PV nuclei and neighboring PNN-negative nuclei (termed non-PV) within S1BF (Figure 1a).

PNN+ PV nuclei of WT exhibited a roughly 3-fold higher level of MECP2 than in non-PV nuclei in Layer IV S1BF across adolescence and adulthood (Figure 1a, b). This fold increase is higher than the 1.25-fold increase reported in Layers II/III (Ito-Ishida et al., 2015). We observed significant increases in MECP2 expression between adolescence and adulthood in WT females across cell types (Figure 1 c, d, f, g). In Het, approximately 50% of the nuclei (identified by DAPI staining) expressed wild-type MECP2 protein at both ages (Figure 1e), in agreement with previous reports (Samaco et al., 2013; Young and Zoghbi, 2004). Strikingly, adolescent Het non-PV nuclei expressed significantly higher levels of MECP2 than age-matched WT, while adult Het non-PV nuclei did not exhibit such a difference (Figure 1c, f). In PV nuclei, MECP2 expression was significantly increased between adolescence and adulthood in both WT and Het (Figure 1d, g), without significant differences between genotypes within ages. Together, these results show that Het displays a precocious increase of wild-type MECP2 levels, as well as an age-dependent failure to further increase wild-type MECP2 expression levels on par with WT females, within specific cell types.

### No changes in perineuronal net expression in S1BF of adolescent Het

MECP2 regulates the expression of perineuronal nets (PNNs), specialized extracellular matrix structures that structurally restrict changes in synaptic plasticity (Carstens et al., 2016, Carstens et al., 2021; de Vivo et al., 2013; de Winter et al., 2016; Deepa et al., 2002; Dityatev et al., 2007; Donato et al., 2013; Durand et al., 2012; Gogolla et al., 2009; Hou et al., 2017; Kosaka and Heizmann, 1989; Krishnan et al., 2015; Krishnan et al., 2017; Morello et al., 2018; Nakagawa et al., 1986; Patrizi et al., 2020; Pizzorusso et al., 2002; Sigal et al., 2019; Thompson et al., 2018; Ueno et al., 2017). Adult Het show atypical increases in the expression of mature, high intensity PNNs in S1BF and the auditory cortex (Krishnan et al., 2017; Lau et al., 2020b). PNNs are formed on PV neurons from an array of secreted proteins sourced from multiple cell types (Bignami et al., 1992; Carulli et al., 2006, Carulli et al., 2007; Kwok et al., 2010; Miyata and Kitagawa, 2017). Thus, we hypothesized that the increased MECP2 levels within non-PV cells would regulate adolescent Het PNN expression to typical WT levels in a non-cell autonomous manner.

As we found nuanced changes in latero-medial axis and a hemisphere-specific difference in PNN expression in adult females (Lau et al., 2020b), we used serial sectioning, immunofluorescent staining, and epifluorescence microscopy to manually quantify high-intensity PNN expression across the entire S1BF (Figure 2a, b). We found no significant difference in high-intensity PNN density between adolescent WT and Het across the entirety of the S1BF (Figure 2c). Additionally, we did not find differences in PNN densities between hemispheres of either adolescent genotype. (WT-L = 12.845 ± 1.607 # PNNs/mm^2^, WT-R = 18.315 ± 1.016, n = 307 images, 6 animals; Het-L = 13.241 ± 0.380, Het-R = 14.932 ± .347, n = 313 images, 6 animals).

Differing intensities of PNNs are thought to represent levels of maturity of that PV network in early developing postnatal cortex. However, it is unclear if the same standards hold true for adolescent cortices. Thus, we performed additional analyses using an automated, unbiased approach to select all pixels that represent PNN expression (Figure 2d, red distributions in top two panels; see Methods) over background (Figure 2d, blue distributions in top two panels). We observed that PNN expression in all tissues had multimodal pixel intensity distributions, which consistently included a bell-shaped distribution at higher intensities (Figure 2d insets). This distribution represents specific cortical layers with strong PNN expression (Layers II/III, IV, and V) of interest for quantification. Models consistently captured the distributions of PNN expression in the expected intensity region (Figure 2d, red). By analyzing the mean distribution of these red traces, we observed no significant differences in PNN signal between adolescent WT and Het using a linear mixed-effects model with cohort as a variable (Figure 2d, bottom panel). The intraclass correlation coefficient across all cohorts was moderate (0.218), demonstrating that variation within the dataset was significantly driven by natural variation due to random X-chromosome inactivation, over animal-to-animal variability (See Materials and Methods). Together, these results suggest that MECP2 expression in non-PV cells affects PNN expression in PV cells in an age-dependent manner.

### Adolescent Het perform a pup retrieval task with comparable efficiency to age-matched WT

Adult Het display sensory deficits while performing the complex social behavior of alloparental care, as measured by the pup retrieval task (Krishnan et al., 2017; Stevenson et al., 2021). Since adolescent Het expressed more MECP2 in non-PV nuclei and expressed PNNs at WT levels, we predicted they would perform efficiently in the pup retrieval task (Figure 3). Adolescent Het and WT improved at pup retrieval across six consecutive days of testing as measured by time to retrieve scattered pups to the nest (latency index; Figure 3a, a’) and number of errors (Figure 3b, b’). Additionally, adolescent Het retrieved pups with a similar efficiency as adolescent WT females. Adult Het are comparably inefficient at this same task (Stevenson et al., 2021). Cumulatively, these results suggest that the increased MECP2 levels within non-PV nuclei, as well as typical PNN expression associated with PV+ interneurons, provide a molecular mechanism for compensatory behavioral adaptations during adolescence which are lost in older females, resulting in a regressive behavioral phenotype.

### Het display regression in whisker tactile sensory behavior between adolescence and adulthood

Pup retrieval involves initial perception of sensory cues, multisensory integration, and motor coordination involving multiple neural circuits. Here, we chose to focus specifically on the single sensory perception modality of whisker tactile perception due its connection to S1BF. We performed standard texture discrimination and object recognition tasks on adolescent and adult WT and Het and analyzed their behavior frame-by-frame using DataVyu software (Figure 4).

Standard representation of percentage of time spent with different textures did not change in adolescent females during texture investigation (WT: 50.14% ± 4.344, n=9: Het: 55.60% ± 4.116, n=10; one-sample t-test with 50% chance) or during object investigation (WT: 47.74% ± 4.263, n=9: Het: 50.33% ± 3.572, n=10; one-sample t-test with 50% chance). Percentage of time spent with textures did not change in adult WT or Het female mice (WT: 57.08% ± 3.356, n=9; Het: 51.59% ± 3.531, n=8; one-sample t-test with 50% chance) and during object investigation (WT: 51.67% ± 4.477, n=9; Het: 57.26% ± 4.511, n=8; one-sample t-test with 50% chance). Thus, we refrain from drawing conclusions about preferences of the female WT and Het mice or their memory capabilities here.

We chose to perform a systematic analysis of behavioral videos, as mice interact with textures and objects with high speed and rarely linger during investigations. We found that both adolescent WT and Het females predominantly investigated textures with whiskers and interacted with similar number of contacts (Figure 5a, b). However, though the total interaction time with textures did not differ significantly between adolescent WT and Het (Figure 5c), the duration per whisker interaction was significantly higher in Het (Figure 5d).

**Figure 5.**
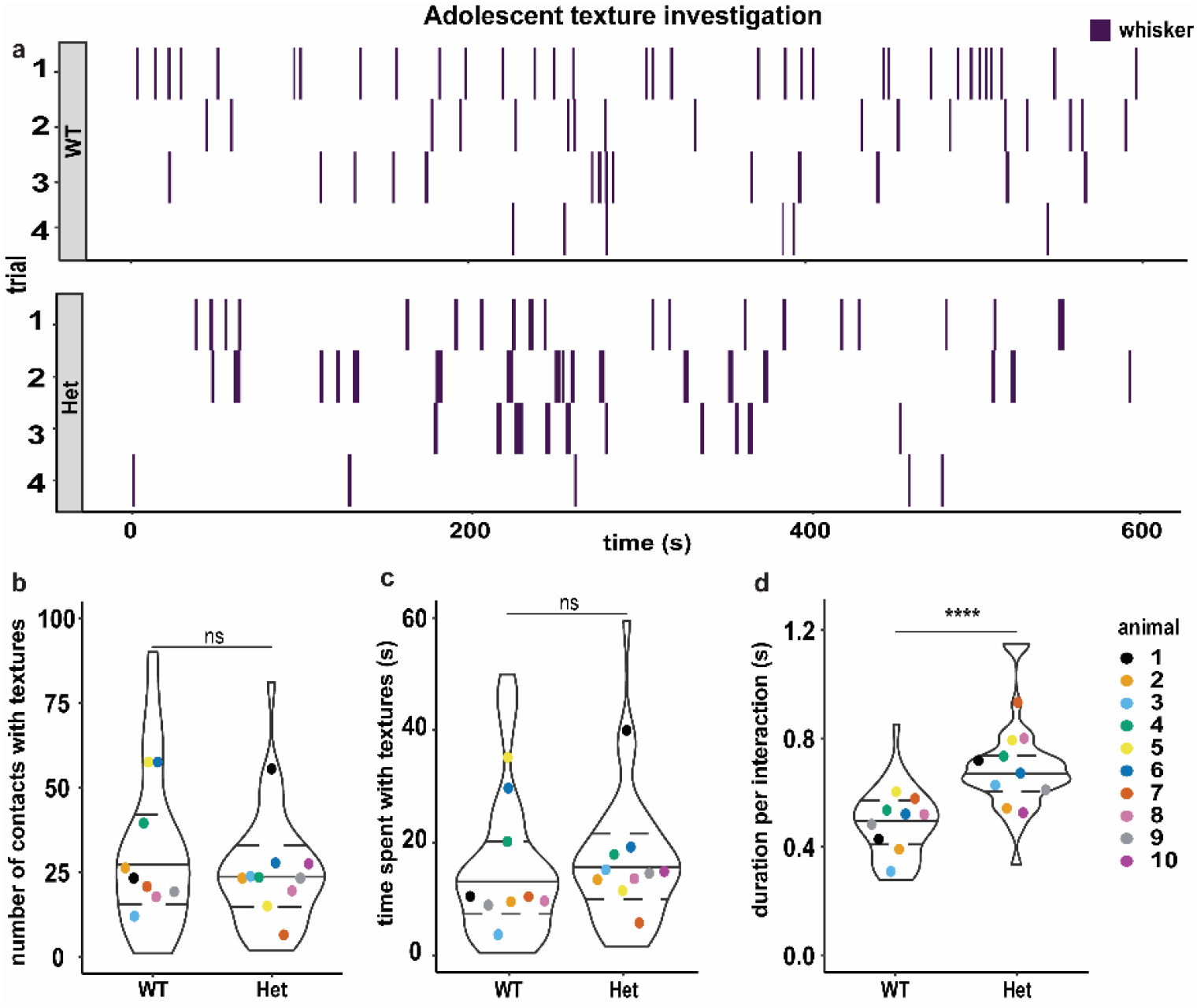
Adolescent Het have subtle whisking deficits during texture interactions. **(a)** Examples of frame-by-frame texture interactions across four 10-min trials of texture exposure for adolescent WT (top panel) and Het (bottom panel), represented as sequences. The X-axis represents the time (s). Whisker interactions (purple) are plotted as blocks that span the duration of the interaction. Each row depicts fleeting whisker interactions of a mouse during individual trials. **(b-d)** Statistical analysis of the interaction quantifications revealed that on average, WT and Het had similar total number of contacts across 2-4 trials (b) and spent similar averaged total time investigating textures across 2-4 trials (c). However, Het spent significantly longer during each whisker interaction than WT (d). Superplots data visualization shows individual cohorts by same color circles, represent the mean of 2-4 trials (n=9 WT, n=10 Het) and violin plots show median and **rlic+T’il’vnfirvn rrf all iorardirvnc** *Mπmi-Whitney test: ns = not significant, ****p < 0.0001.*

We performed similar analyses with novel object recognition to determine if adolescent Het are deficient in other forms of tactile sensory-dependent tasks. Adolescent WT and Het used their whiskers, forepaws, and whole-body climbing to investigate objects (Figure 6a). Interestingly, adolescent Het made significantly fewer whisker contacts (Figure 6b) and spent similar total time investigating objects (Figure 6e) compared to age-matched WT. There was no significant difference in the number of contacts or time spent investigating objects using forepaws (Figure 6c, f) and whole-body climbing (Figure 6d, g). Similar to texture investigation, adolescent Het interacted significantly longer per interaction using whiskers (Figure 6h), but not forepaws or whole-body climbing (Figure 6i, j), compared to WT. Together, the increased duration of interactions when interacting with textures and objects suggests the beginnings of an atypical whisker perception in adolescent Het.

**Figure 6.**
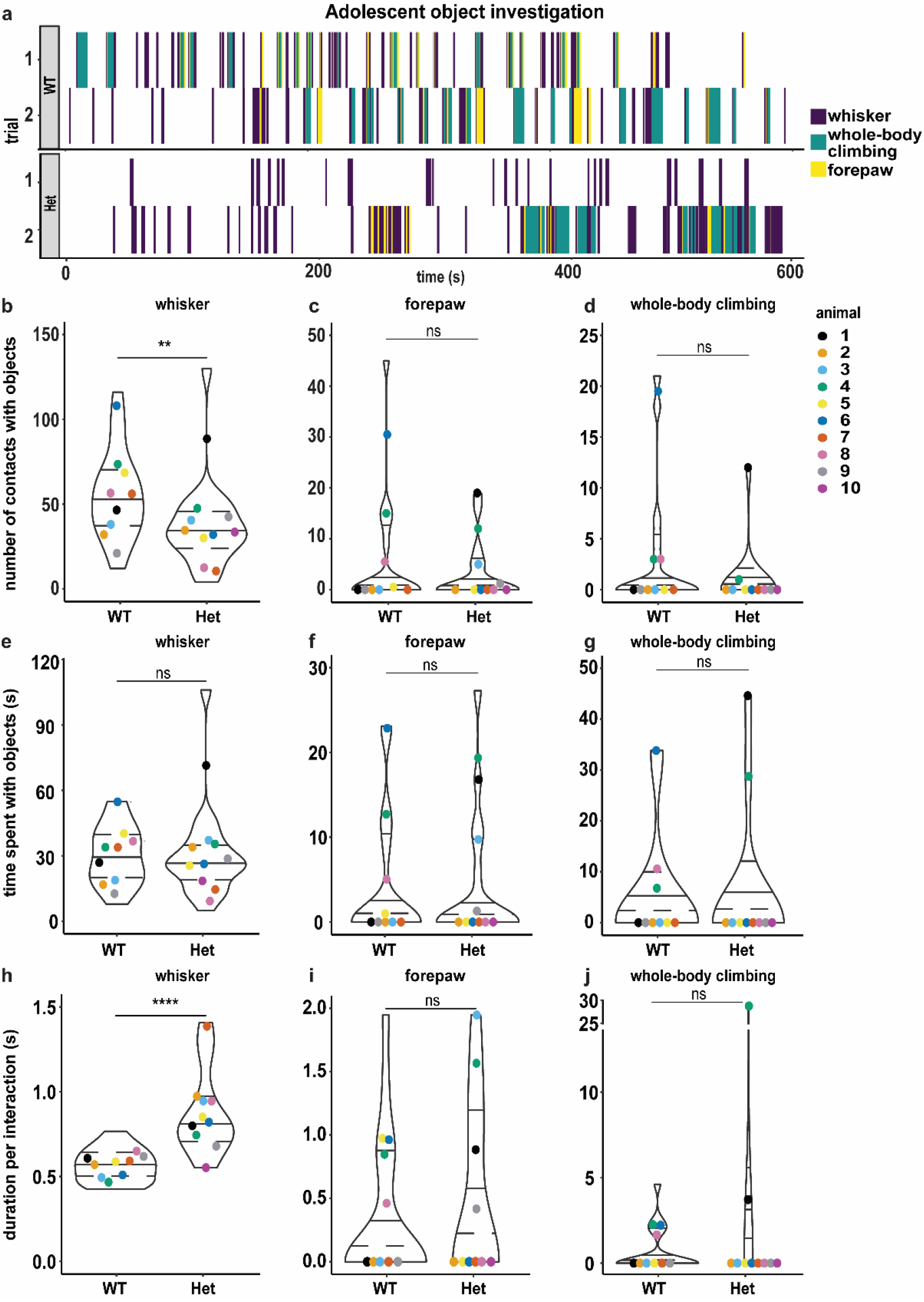
Adolescent Het exhibit subtle whisking deficits during object interactions. **(a)** Examples of frame-by-frame texture interactions across two 10-min trials of object exposure for adolescent WT (top panel) and Het (bottom panel) color coded by types of interaction (whisker, forepaw, whole-body climbing), represented as sequences. The X-axis represents the time (s), and interactions are plotted as blocks that span the duration of the interaction. Each row depicts fleeting whisker (purple), forepaw (yellow) or whole-body climbing (teal) interactions of the mice during individual trials. **(b-d)** Statistical analysis of the interaction quantifications revealed Het made significantly more contacts using whiskers (b), but not forepaws (c) or whole-body climbing (d) compared to WT. **(e-g)** There were no statistical differences between WT and Het for the time spent with objects using whiskers (e), forepaws (f), or whole-body climbing (g). **(h-j)** Het spent significantly more time than WT during each whisker interaction (h), but not with forepaw (i) or whole-body climbing (j). **(b-j)** Superplots data visualization shows individual cohorts by same color circles, represent the mean of 2 trials (n=9 WT, n=10 Het) and violin plots show median and distribution of all interactions. *Mann-Whitney test: ns = not significant, **p < 0.01, ****p < 0.0001.*

On the contrary, adult Het displayed significant whisker hyposensitivity in texture discrimination compared to adult WT (Figure 7). Examples of frame-by-frame sequence analysis showed that non-whisker interactions (forepaw and whole-body climbing) were interspersed with whisker interactions in adult

**Figure 7.**
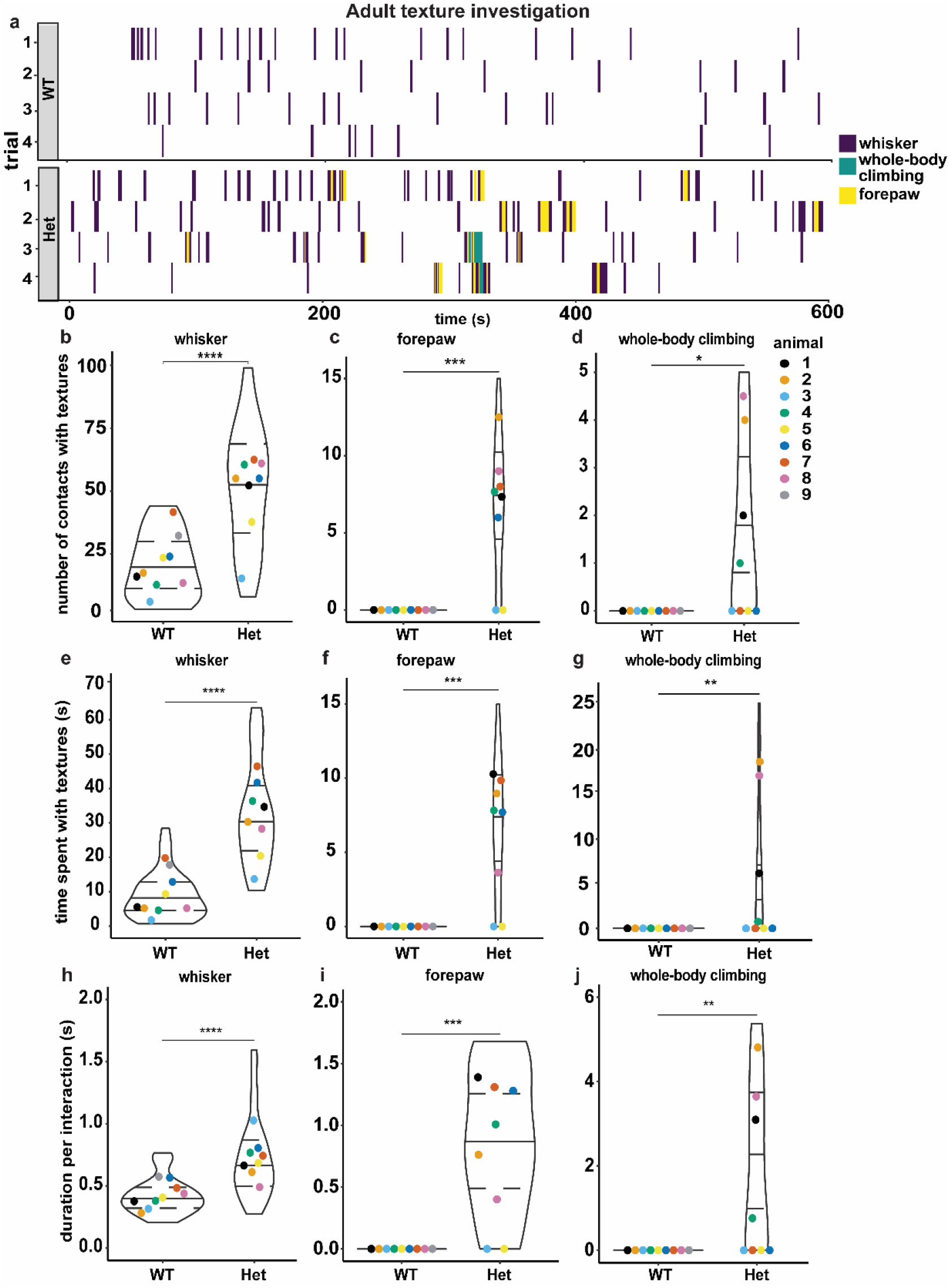
Adult Het exhibit hyposenstivity and compensatory behaviors during texture interactions. **(a)** Examples of frame-byframe texture interactions across four 10-min trials of texture exposure for adult WT (top panel) and Het (bottom panel) color coded by types of interaction (whisker, forepaw, whole-body climbing), represented as sequences. The X-axis represents the time (s), and interactions are plotted as blocks that span the duration of the interaction. Each row depicts fleeting whisker (purple), forepaw (yellow) or whole-body climbing (teal) interactions of the mice during individual trials. Statistical analysis of the interaction quantifications revealed adult Het made significantly more contacts (b-d), spent significantly more total time (e-g) and time per interaction (h-j) with textures, with all interaction types (b-j), compared to adult WT. Superplots data visualization shows individual cohorts by same color circles,, represent the mean of 2-4 trials (n=9 WT, n=8 Het) andviolin plots show median and distibution of all interactions. *Mann-Whitney test: *p < 0.05, **p < 0.01, ***p < 0.001, ****p < 0.0001.*

Het (Figure 7a). Compared to WT, adult Het used whisker to make significantly more contacts, spent more time investigating textures, and had longer durations per whisker interaction with textures (Figure 7b, e, h). Adult Het also used forepaws and whole-body climbing to make significantly more contacts, spent more time investigating, and had longer time per interaction with textures than adult WT (Figure 7c-d, f-g, i-j, respectively). These findings are similar to previous studies in whiskerless and barrel-less mice (Guic-Robles et al., 1992; Guić-Robles et al., 1989) and suggest adult Het are seeking compensatory tactile sensation due to deficits in whisker sensation (hyposensitivity). Furthermore, this systematic analysis captured the individual variation expected in adult Het phenotypes. Adult Het 3 and 5 (Figure 7b-d, h-j, light blue and yellow colors respectively) were similar to adult WT in number and duration of contacts with whiskers, and no interactions with forepaw or whole-body, suggesting a likely typical functionality within S1BF of the respective animals. Adult Het 6 and 7 (dark blue and deep orange respectively) performed forepaw interactions, but not whole-body climbing (Figure 7c, d, f, g, i, j), suggesting minor sensory processing deficits in S1BF which might be compensated for by appropriate tactile perception within the forelimb region of the primary somatosensory cortex. The other five adult Het were most different from adult WT in their texture interactions, likely to have profound deficits in neural processing in S1BF and neighboring SS1 subregions.

Surprisingly, adult Het had milder phenotypes while interacting with objects (Figure 8). Unlike texture investigations in adults, both adult WT and Het also used forepaws and whole-body climbing to interact with objects (Figure 8a). We did not observe significant differences in both the total number of contacts with objects and the total time spent investigating objects between adult WT and Het during whisker, forepaw, or whole-body climbing interactions (Figure 8b-g). Adult Het spent significantly longer time per interaction with objects using whiskers (Figure 8h), but not forepaws or whole-body climbing (Figure 8i, j). Taken together, these results point to the beginnings of a regressive phenotype associated with specific tactile sensory perception in 6-week-old female Het that progresses to a severe, context-dependent tactile hyposensitivity by 12 weeks of age.

**Figure 8.**
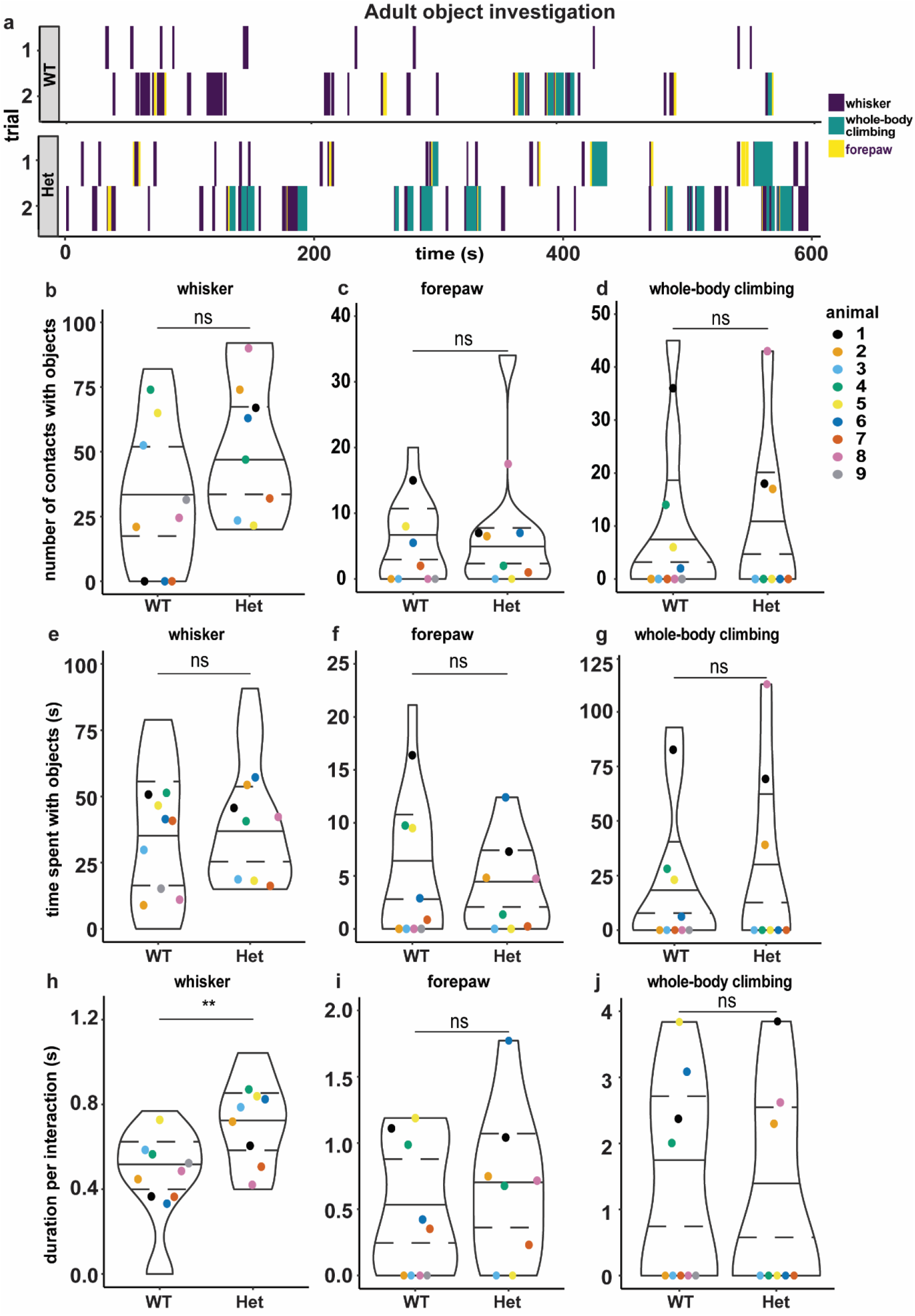
Adult Het display mild whisker deficit while interacting with objects. **(a)** Examples of frame-by-frame texture interactions across two 10-min trials of object exposure for adult WT (top panel) and Het (bottom panel) color coded by types of interaction (whisker, forepaw, whole-body climbing), represented as sequences. The X-axis represents the time (s), and interactions are plotted as blocks that span the duration of the interaction. Each row depicts fleeting whisker (purple), forepaw (yellow) or whole-body climbing (teal) interactions of the mice during individual trials. **(b-j)** Statistical analysis of the interaction quantifications revealed adult Het interacted with objects using whisker, forepaw and whole-body climbing, similar to WT, in the number of contacts (b-d), total time with objects (e-g) and time per interactions (with Het having a slight but significant increase with whisker) (h-j). Superplots show animals color coded by circles, represent the mean of 2 trials (n=9 WT, n=8 Het). Superimposition of violin plots show median and distribution of all interactions. Statistical significance determined by *Mann-Whitney test: **p < 0.01.*

### Context-specific developmental progression in WT and regression in Het

To develop a baseline in tactile performance of females in both adolescence and adulthood, we compared texture- and object-associated phenotypes within genotype and across age. Compared to adolescent WT, we found that adult WT spent significantly lesser time with textures (Figure 9a, b), while the number of contacts remained similar across ages (Figure 9c). On the other hand, adult Het spent significantly more time with textures and more numbers of contacts (Figure 9d, f) compared to adolescent Het, while the duration per contact remained similar (Figure 9e). Together, these results suggest a progressive perception via whisker tactile interactions over age in WT, while Het display regressive inefficiencies by these same parameters. Interestingly, neither WT nor Het showed any significant age-specific changes in interactions with objects (Figure 10a-f), suggesting that female mice may use different strategies while interacting with finer textures and objects.

**Figure 9.**
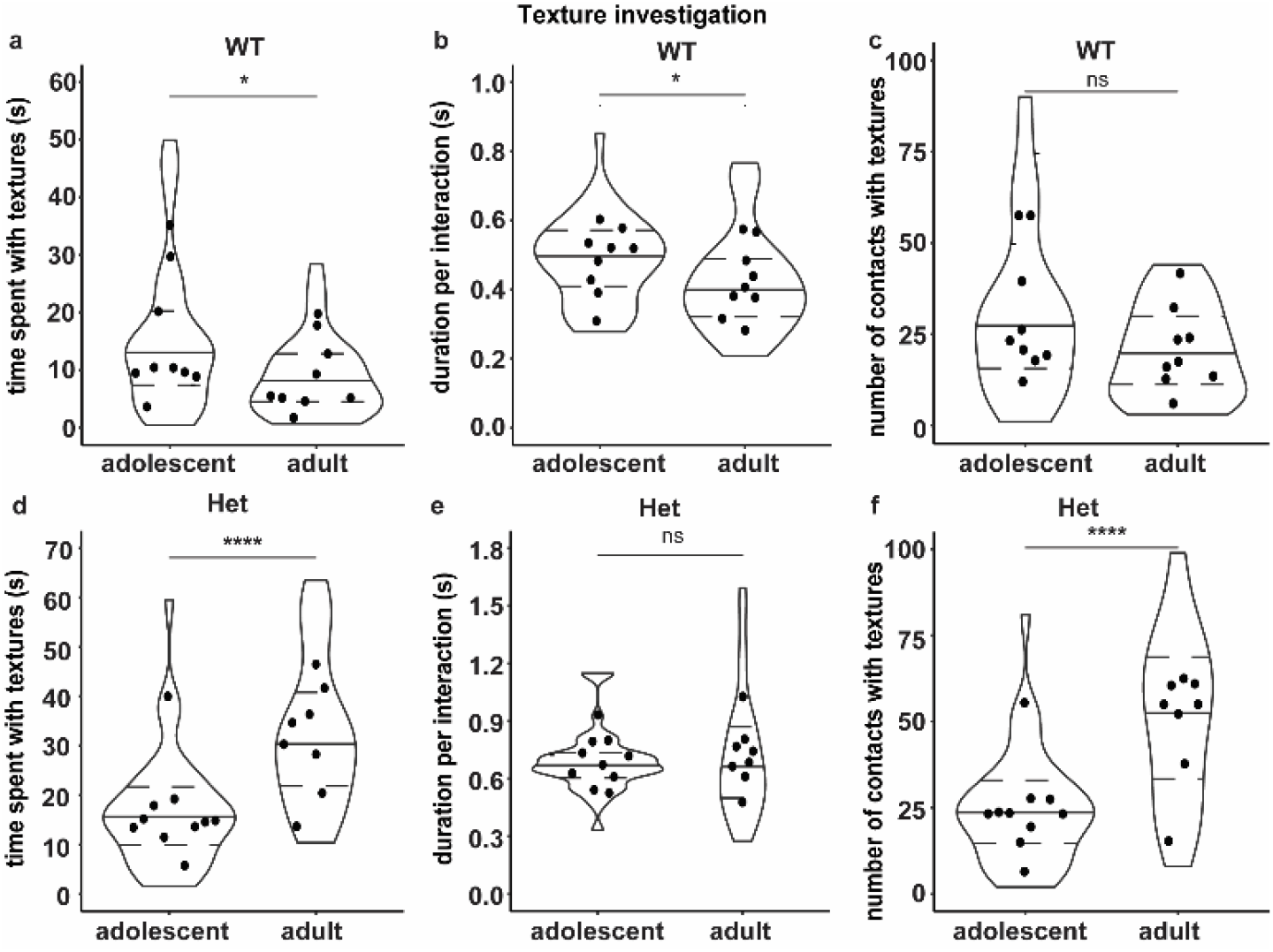
Differential and agespecific changes in WT and Het during texture interaction. **(a-c)** Using whiskers, adult WT interacted in significantly shorter time (a: total time, b: time per interaction) with textures than adolescent WT, without changes in number of contacts made (c). **(d-f)** However, adults Het used whiskers to significantly spent longer time (d) and made more contacts (f) with textures than adolescent Het, without changes in time spent per interaction (e). **(a-f)** Each circle represents the mean of 2-4 trials per animal (adolescent: n=9 WT, n=10 Het; adult: n=9 WT, n=8 Het). Superimposition of violin plots show median and distribution of all interactions. *Mann-Whitney test: ns = not significant, *p < 0.05, ****p < 0.0001.*

**Figure 10.**
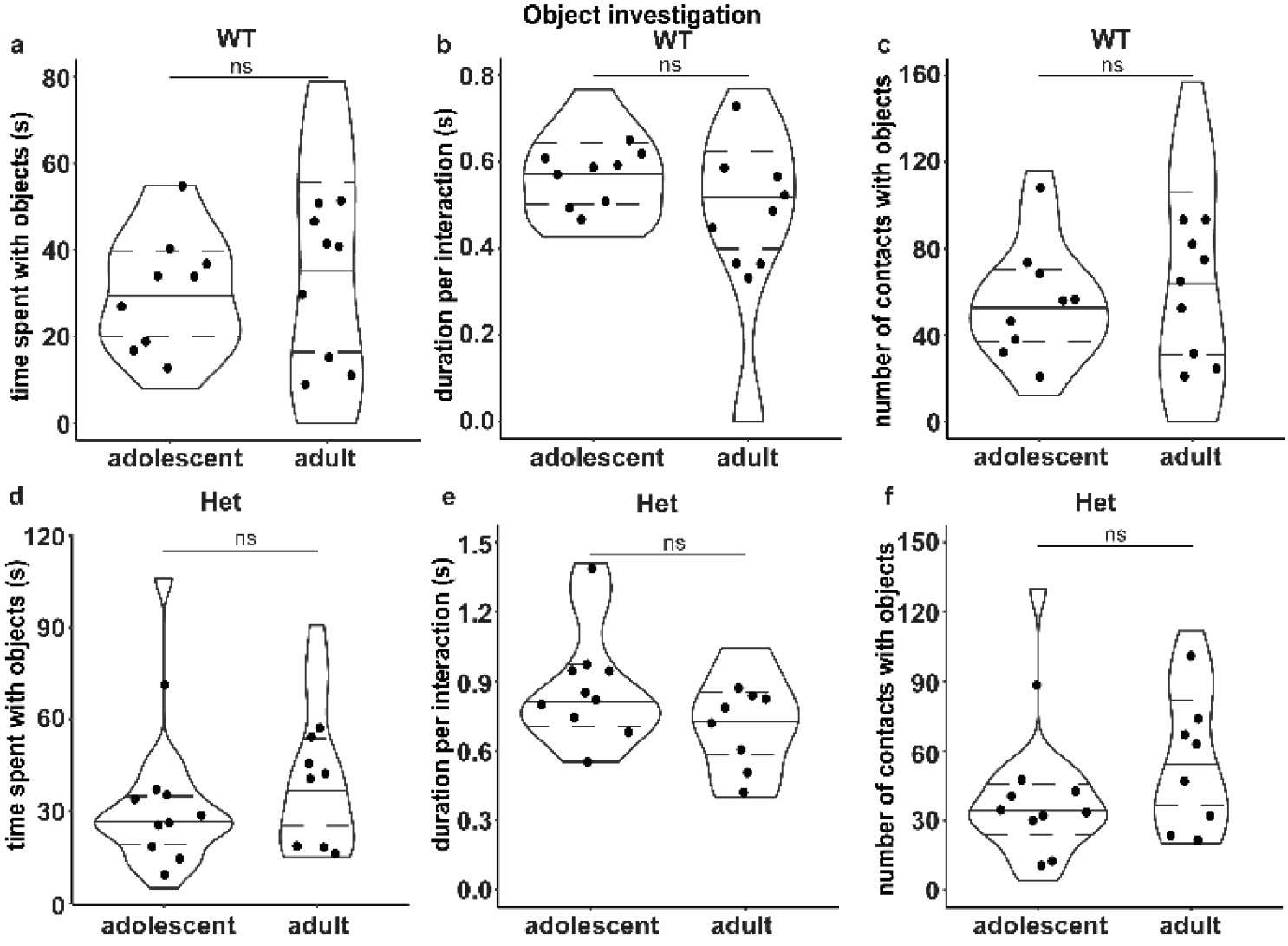
No differential or agespecific changes in WT and Het during object interaction. **(a-c)** Both adolescent and adult WT used whiskers similarly to interact with objects, as determined by total time spent (a), time per interaction (b) and number of contacts (c). **(d-f)** Comparably, adolescent and adult Het used whiskers similarly to interact with objects (d: total time; e: time per interaction; f: number of contacts). Each circle represents the mean of 2 trials per animal (adolescent: n=9 WT, n=10 Het; adult: n=9 WT, n=8 Het). Superimposition of violin plots show median and distribution of interactions. *Mann-Whitney test: ns = not significant.*

## Discussion

Disrupted sensory processing is a contributing factor to many of the behavioral phenotypes in neurodevelopmental disorders (Rais et al., 2018; Schaffler et al., 2022). Here, we use established models of whisker tactile sensory perception to discover the impact of mosaic MECP2 expression in identified cell types across developmental stages in a female mouse model for Rett syndrome (Figure 11). These results set the stage to explore the mechanistic links between age-dependent MECP2 expression levels to neural circuits involved in sensory perception during simple and complex behavioral tasks within Het. Two major observations are of importance to the RTT field: (1) wild-type MECP2 protein expression is regulated in an age- and cell-type specific manner in Het, providing an unexplored angle for therapeutic intervention in the Het brain via dynamic protein regulation, and (2) female Het mice have typical functioning capacity which is lost with age, suggesting that the Het brain is capable of cellular and behavioral plasticity, even past early infancy. Deep phenotyping at cellular and behavioral levels is necessary in this important preclinical model of Rett syndrome to aid in better therapeutic treatments.

**Figure 11.**
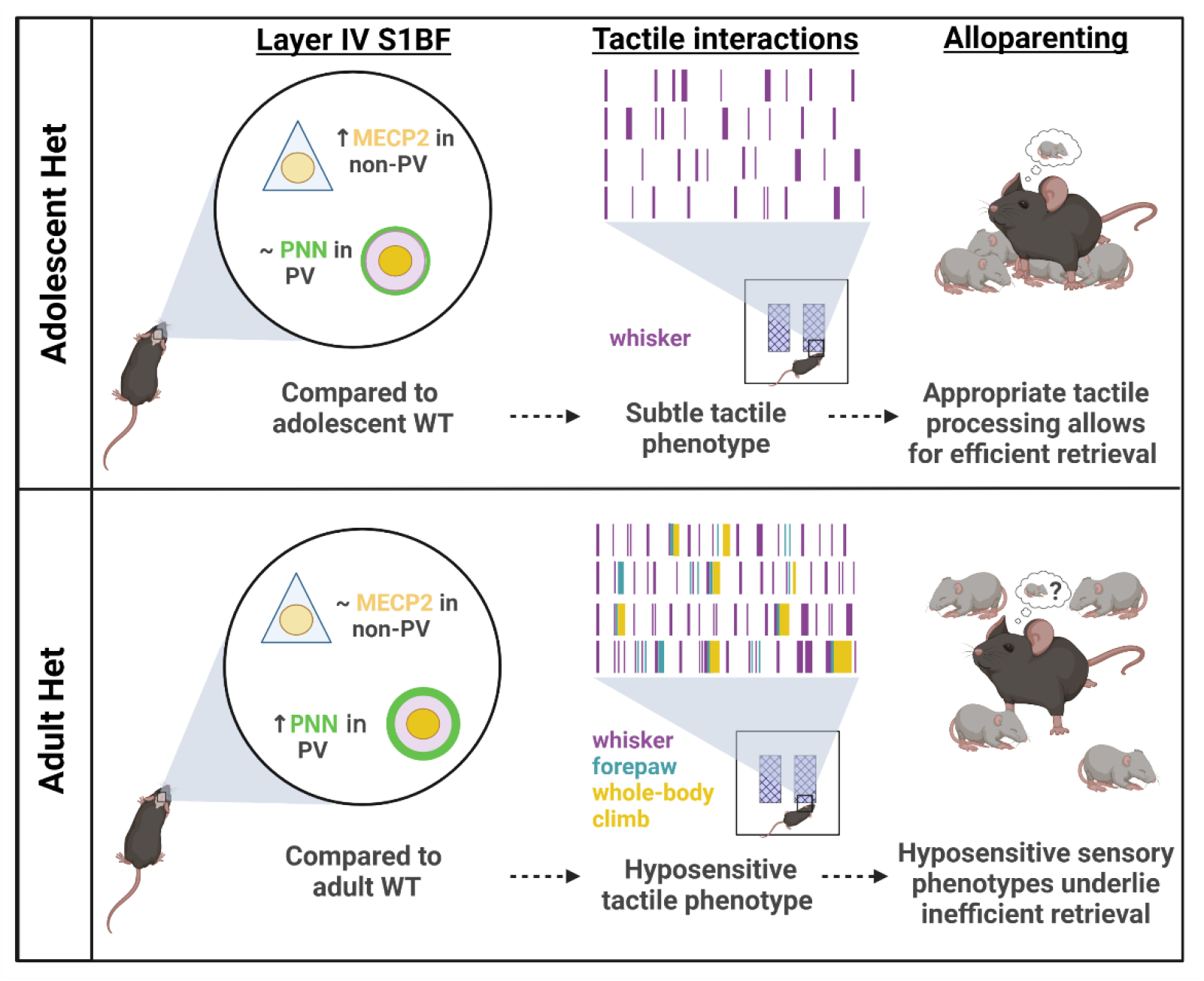
Summary diagram: MECP2 and PNN expression coincide with age-dependent sensory deficits in Het. Wild-type MECP2 expression is dysregulated in genotypic adolescent *Mecp2*-heterozygous female mice, which coincides with typical expression of perineuronal nets in barrel field of the primary somatosensory cortex, mild tactile sensory perception, and efficient multisensory processing during the pup retrieval task. In adulthood, wild-type MECP2 expression in Het is similar to control WT brains, which coincides with atypical abnormal increase in perineuronal nets, strong whisker interactions indicative of a hyposensitivity phenotype, and aberrant multisensory processing leading to inefficient pup retrieval.

### Aberrant homeostasis of MECP2 in late adolescence as a catalyst for the transition from pre-symptomatic to symptomatic RTT female mice

One of the most intriguing results here is the observed differences in MECP2 expression within nuclei of Layer IV S1BF non-PV cells in adolescent Het compared to age-matched WT. The observation of cell-type specific and age-dependent MECP2 expression dynamics within Het provides new avenues for consideration in RTT pathogenesis. To date, Rett syndrome is thought to be caused by the lack of functional MECP2 in roughly half of all cells. We speculate that the transient fluctuation in MECP2 levels within the ~50% of the cells expressing the wild-type protein is an additional direction to consider. The precocious increase in MECP2 within non-PV nuclei of Het may be a cell-autonomous mode of compensation for the lack of MECP2 protein in surrounding cells. Over time, however, the ability to further increase MECP2 expression levels in non-PV nuclei, as observed in all WT cells as well as Het PV cells, is lost.

Thus, although the genotype and the cellular status of the Het remain the same throughout life, their MECP2 expression levels are changing in a cell-type specific manner, suggesting built-in homeostatic mechanisms to regulate MECP2 expression. This could be a basis for normal development in infancy for girls diagnosed with RTT, and then regression at different times throughout life (Neul, 2019). It is equally intriguing that MECP2 expression in PV neurons, with higher basal levels than in non-PV cells, exhibit typical increases comparable to WT females across age. Yet, the underlying mechanisms regulating MECP2 expression levels within specific cell types across age remain unclear. Sorting specific cell types and measuring MECP2 protein content, with regional and age specificities, are necessary to establish an extensive expression baseline in Het, as well as to determine how MECP2 expression changes with experience within context-specific circuitry.

In female Het mice, the precocious, cell-type specific increases in MECP2 expression coincides with pubertal changes. In female WT mice, onset of puberty is accelerated by leptin (Ahima et al., 1997; Chehab et al., 1997) and ovarian hormones that regulate the organization and maturation of cortices by regulating inhibition (Clemens et al., 2019; Piekarski et al., 2017). This line of research is important as girls with RTT enter puberty early and reach menarche later, and body-mass index affects pubertal timing (Killian et al., 2014). If Het mice also undergo pubertal deviations from their WT littermates, atypical neuroendocrine signaling would affect Het females as they transition into adulthood, in terms of both steroid-dependent organization of circuitry and MECP2 expression.

MECP2 expression changes in appropriate primary sensory cortices in early developmental critical periods (He et al., 2014; Krishnan et al., 2015; Skene et al., 2010; Yagasaki et al., 2018) and in response to different stimuli (Martínez de Paz et al., 2019; Yagasaki et al., 2018). MECP2 expression in adolescence and adulthood is thought to be stable. However, systematic studies with celltype specific fluorescent-assisted cell sorting over brain regions, ages, and experimental conditions are required to determine the extent of the dynamic range of MECP2 expression. Does the expression level of MECP2 matter for its nuclear functions of binding methylated DNA and regulating gene expression? Though not thoroughly investigated at the mechanistic level, increasing MECP2 expression in transgenic over-expression rodent lines (Collins et al., 2004) or deleting MECP2 in specific cell types also largely results in Rett syndrome-like phenotypes at behavioral and cellular levels (Bhattacherjee et al., 2017; Chapleau et al., 2009; Goffin et al., 2014; He et al., 2014; Ito-Ishida et al., 2015; Krishnan et al., 2017; Meng et al., 2016; Mossner et al., 2020). However, how MECP2 expression in neighboring cells are affected is unknown in these manipulations. Moreover, considering the possibility of compensatory changes in MECP2 expression, it is critical to determine the impact of these manipulations on MECP2 expression in unmanipulated, non-target cells in an age-specific manner. Furthermore, we speculate such compensatory mechanisms at the protein level to play crucial roles in other X-linked neurodevelopmental disorders (Brand et al., 2021; Santos-Rebouças et al., 2020) and metabolic disorders (Juchniewicz et al., 2021).

### Perineuronal nets and PV-pyramidal networks in *Mecps*-deficient mouse models

Germline deletion of *Mecp2* from all cell types, as well as conditional deletion of *Mecp2* from both PV neurons and forebrain pyramidal neurons, result in myriad of cellular and behavioral phenotypes, particularly in male mice (Chao et al., 2010; Gemelli et al., 2006; Ito-Ishida et al., 2015; Krishnan et al., 2015; Morello et al., 2018; Mossner et al., 2020; Patrizi et al., 2020). An altered PV-network configuration interferes with experience-dependent plasticity mechanism in the brain and is associated with defective learning and plasticity (Donato et al., 2013; Krishnan et al., 2015; Patrizi et al., 2020). Adult Het mice have atypical increased expression of PNNs in the auditory cortex (Krishnan et al., 2017) and specific subregions of the primary somatosensory cortex (Lau et al., 2020b). Typically, pup vocalizations trigger suppression of PV+ GABAergic neuronal responses and a concomitant disinhibition in deep-layer pyramidal neurons in the auditory cortex of adult WT surrogates (Lau et al., 2020a). In adult Het surrogates, the increased perineuronal net expression interferes with the synaptic plasticity of PV+ GABAergic neurons, resulting in a lack of disinhibition of pyramidal neurons (Lau et al., 2020a) and coinciding with poor pup retrieval performance (Krishnan et al., 2017). It is currently unknown if similar mechanisms also occur in the primary somatosensory cortex, and if so, whether they occur in an age-dependent manner, which would ultimately lead to atypical tactile processing in Het. Studies are underway to determine how PNN expression in S1BF contributes to tactile sensory perception.

### Systematic analysis of behavioral assays to study tactile sensory perception in female mice

Traditionally, texture discrimination and object recognition assays are reported as “percent novelty recognition” or a “discrimination index” calculated as total time spent investigating the novel texture/object divided by the total time spent investigating both textures/objects (Antunes and Biala, 2012; Ennaceur et al., 2005; Ennaceur and Delacour, 1988). This analysis normalizes the time spent investigating across individual animals to control for variation in exploration time. Based on these metrics, a mixed result of tactile hyper- and hypo-sensitivity has been previously reported in models of MECP2 deficiency (Chao et al., 2010; Ito-Ishida et al., 2015; Orefice et al., 2016; Chen et al., 2001; Gemelli et al., 2006). It is hypothesized that mice will spend more time with novel texture/object due to their innate nature of curiosity and their ability to remember old or novel objects/textures from such interactions. However, a growing body of literature shows that free-moving rodents do not have a preference for novelty (Antunes and Biala, 2012; Cohen and Stackman Jr., 2015; Ennaceur, 2010). Mice and rats show preference for familiar objects rather than novel objects (Ennaceur, 2010). Adult C57BL/6 male mice show novelty object preference, whereas C57BL/6 adult females show no preference between a familiar and novel object (Frick and Gresack, 2003), consistent with our findings. Thus, to reconcile these differences in the literature and establish first order metrics for analysis, we performed a systematic frame-by-frame analysis of interactions with textures and objects for adolescent and adult female C57BL/6 mice. By analyzing the number and time of interactions, we found that adult WT females spent less time interacting with textures compared to adolescent WT females. On the other hand, adult Het females interacted more overall and longer than adolescent Het females, suggesting a regressive tactile hyposensitivity phenotype in adult Het.

### Regression in RTT models

Previous work using a female rat RTT model has reported a regression in a learned forepaw skill over the course of weeks (Veeraragavan et al., 2016). Regressive breathing abnormalities have also been shown using a conditional MECP2 knockout male mouse model (Huang et al., 2016). Previously, we have proposed that adult female Het show regression in maintaining pup retrieval efficiency over few days (Stevenson et al., 2021). While these studies report regression in parasympathetic and motor function, our work here is the first to report a regressive hyposensitivity phenotype in tactile perception between adolescence and adulthood in a female mouse model for RTT. An alternative explanation for increases in tactile interactions from adolescence to adulthood in Het is the emergence of repetitive obsessive-compulsive disorder-like phenotypes over time, as observed in mouse models of this disorder (Ahmari, 2016; Alonso et al., 2015; Mitra and Bult-Ito, 2021). Adult Het perform repetitive jumping during early days of pup retrieval which could be indicative of novelty- or stress-induced stereotypies in a subpopulation of adult Het (Stevenson et al., 2021). It is currently unclear if the increased frequency and longer durations of interactions in adult Het are a form of repetitive obsessive-compulsive behavior. Clozapine injection has been shown to exacerbate such repetitive phenotypes (Kang et al., 2020) and can be utilized to test this hypothesis in the future.

### Rehabilitation strategies

Early interventions aimed at increasing neuronal activity mitigate disease onset in both male and female mouse models for Rett syndrome. Early administration of Ampakine CX546, a positive modulator of AMPA receptors, from postnatal days 3-9 in male *Mecp2*-null mice enhances neuronal activity, prolongs lifespan, delays disease progression, and rescues both motor abilities and spatial memory (Scaramuzza et al., 2021). In Het females, repetitive motor training enhances neuronal activity of task-specific neurons and improves both spatial memory and motor function well into adulthood (Achilly et al., 2021). Adolescent and adult Het improve in the pup retrieval task with repeated trials over days, in contrast to non-consecutive days of training (Krishnan et al., 2017; Stevenson et al., 2021). Early and repetitive trainings delay the onset of disease phenotypes and improve the latency to survival in Het, supporting the plastic capacity of the adolescent Het brain (Achilly et al., 2021). We speculate that wild-type MECP2 expression in Het is affected by these trainings, especially at earlier ages, resulting in activation of compensatory pathways which lead to appropriate behavioral adaptations. Further work is needed to identify therapeutic and rehabilitative strategies that target MECP2 maintenance within the female Het brain for mitigating disease progression throughout life.

## Acknowledgements

We would like to thank undergraduates Beyza Kartal, Benjamin Alireza Bridges, Sami Issac, Shahin Ahmadi, Itzanami Sotelo Hernandez, Miranda Blevins and Deema Mansour for technical help, and the 2020 and 2021 BCMB 420 Image analysis class for help in mapping and analyzing high intensity PNN data. We would like to thank the lab of Dr. Maitreyi Das and her graduate students for training and assistance using the confocal microscope. This work was supported by BCMB Chancellor’s fellowships (MM and DLC), the 2021 Scholarly and Research Incentive Funds (SARIF) through the Office of Research, Innovation, & Economic Development (ORIED) at UTK (MM), BCMB James and Dora Wright Fellowship (MM), W. Sherman Kouns Excellence in Teaching award (MM), National Science Foundation Graduate Research Fellowship Program (GRFP) (LD), postdoctoral fellowship award from Rettsyndrome.org (BYBL), startup funds from the University of Tennessee—Knoxville (KK), and by the National Institute of Mental Health of the National Institutes of Health under Award Number R15MH124042 (KK).

## Contributions

KK and DLC designed the study. DLC and DWS performed tactile behavior. LD, MM, TRS and BYBL performed 6-week-old consecutive maternal behavior. TRS and AM scored maternal behavior and BYBL checked scores.

LD performed confocal imaging and MECP2 expression analysis. DLC, AM and SP performed data quantification in Datavyu. DLC performed immunohistochemistry and epifluorescent PNN imaging for 6-week-old brains. DLC, AM and SP performed mapping and PNN high intensity count analysis for 6wk naïve animals. LD and SP collected region-of-interest histogram data for all-intensity PNN analysis. MM designed the R studio codes for analysis and data visualization of tactile behavioral data. LD performed statistical analysis for MECP2 expression and generated figures in GraphPad. AW and TH performed analysis of PNN histogram data. TRS performed genotyping of mice. MM, LD, DLC, BYBL and KK contributed to in-depth discussions, and wrote the paper.

